# Dietary tryptophan rescues growth impairment during giardiasis independently of parasite burden

**DOI:** 10.64898/2026.10.29.762798

**Authors:** Rita Kosile, Vanessa Angelova, Peiran Shen, Bergeline C. Nguemwo-Tentokam, Leila Pagel, Evan Pannkuk, Luther Bartelt, Steven M. Singer

## Abstract

Giardiasis is an intestinal disease caused by the parasite *Giardia duodenalis*. Intestinal barrier dysfunction during *Giardia* infection has been linked to childhood growth stunting, particularly in communities with prevalent malnutrition and repeated exposure to *Giardia*. In pediatric cohort studies, growth stunting has been strongly associated with asymptomatic *Giardia* detection in fecal samples. Several studies have described potential mechanisms by which *Giardia* disrupts the epithelial barrier, but less is known about factors that may support epithelial homeostasis in response to infection. Microbiota-derived tryptophan metabolites that activate aryl hydrocarbon receptor (AHR) signaling are a key pathway involved in epithelial barrier homeostasis. We tested whether dietary tryptophan availability influenced host growth outcomes and aryl hydrocarbon receptor activity during giardiasis. We used established murine *Giardia* malnutrition models and compared host growth, a correlate of barrier dysfunction, with a tryptophan-supplemented diet group. *Giardia*-infected mice fed a low-protein diet failed to gain weight whereas infected mice fed an isocaloric regular protein diet gained weight similar to diet-matched controls. Tryptophan supplementation of the low-protein diet rescued this diet-dependent and infection-induced weight defect in C57BL/6J and BALB/c mice. This growth rescue was associated with increased intestinal Cyp1a1 expression, a canonical AHR target, but without a reduction in infection burden. Glycine supplementation, used as a non-essential amino acid control that is not expected to generate AHR ligands, did not produce the same effect, suggesting that the growth rescue observed with tryptophan supplementation was not due to general amino acid supplementation. Measurements of circulating tryptophan metabolites in plasma showed limited and inconsistent differences across experimental groups, which may suggest that the growth rescue phenotype may be dependent on local intestinal or mucosal AHR-associated signaling rather than an increase in circulating metabolites. Our findings show that dietary tryptophan availability shapes host growth outcomes during giardiasis. This may also be important as it highlights that host nutrition may be a key contributor to post-infection growth impairment in young children.

**AUTHOR SUMMARY:** *Giardia duodenalis* is a common intestinal parasite that causes intestinal disease extending beyond diarrhea. In children under two years of age, especially in low-and middle-income countries where malnutrition and frequent exposure to *Giardia* is common, *Giardia* infection is often asymptomatic and associated with prolonged small intestinal barrier disruption and decreased length-for-age Z-scores. We wanted to determine whether targeted nutritional supplementation supports resilience to *Giardia* infection. We focused on tryptophan, an essential amino acid that the host and gut microbiota can metabolize into indoles that help regulate intestinal homeostasis. Since *Giardia* can alter the intestinal environment and microbiota, we asked whether dietary tryptophan availability influences host growth outcome. Using mouse models of giardiasis, we found that infected mice fed a low-protein diet failed to gain weight compared with those on a regular protein diet. Adding tryptophan to the low-protein diet improved weight gain in these animals, and this effect was independent of parasite burden. This suggests that tryptophan supplementation may influence host infection tolerance or recovery without accelerating clearance. Tryptophan supplementation also increased expression of an intestinal AHR signaling marker that is key to intestinal barrier function. Our findings suggest that specific dietary alterations may influence post-infection recovery and how malnutrition may contribute to the vulnerability of certain individuals to the long-term effects of giardiasis.

---

*Giardia duodenalis* is a common intestinal protozoan parasite transmitted primarily through ingestion of contaminated food or water [1]. There are an estimated 280 million symptomatic cases annually worldwide. Although symptomatic giardiasis commonly presents with diarrhea, bloating, and abdominal pain, most infections in children with endemic exposures are asymptomatic,. [2]. Even in the absence of diarrhea, large cohort studies, such as Malnutrition and Enteric Disease (MAL-ED) study and the Global Enteric Multicenter Study (GEMS), have consistently associated the presence of *Giardia* with childhood growth stunting [3,4]. *Giardia* however, does associate with increased intestinal permeability [3,5–7]. These findings indicate that childhood growth impairment during giardiasis may result from subclinical or ongoing intestinal barrier dysfunction.

Experimental models have identified several putative mechanisms that may contribute to childhood growth faltering due to *Giardia* infection. Among these, one of the most consistent features of *Giardia-*mediated pathology across epithelial cell culture systems, animal models, and human studies is the disruption of intestinal epithelial barrier integrity [7–13]. *Giardia* trophozoites directly increase intestinal epithelial cell permeability and alter tight junction organization, which can compromise electrochemical gradients and impair nutrient absorption [10,11,14,15]. As with diarrheal symptoms, malnourished children with *Giardia* typically lack a robust mucosal inflammatory response, or immune-associated pathologies like villus atrophy and crypt hyperplasia [16]. Rather, findings from these studies suggest persistent barrier disruption provides a more plausible mechanism linking *Giardia* infection to growth faltering in children in the absence of overt inflammation and diarrhea.

Prolonged developmental sequelae undernourished children with *Giardia* infection suggests an inability to repair the injured epithelial barrier. An injury to the intestinal barrier requires rapid epithelial restitution, followed by the proliferation and differentiation of stem cells in the crypt base to restore epithelial structure and function [17–19]. Evidence suggesting barrier repair post-*Giardia* infection in murine models is the dynamic alterations in cytoskeletal regulatory proteins during infection and resolution. Ezrin and villin are actin-associated cytoskeletal proteins that play critical roles in maintaining epithelial structure and barrier function. Early after *Giardia* infection, intestinal epithelial cells exhibit reduced phosphorylation of ezrin, accompanied by increased phosphorylation of villin. As infection resolves, an accumulation of cleaved ezrin products is observed, suggesting regulated proteolysis of these cytoskeletal proteins during epithelial recovery [20,21]. Consistent with repair as a beneficial response, in a murine model of juvenile *Giardia* (H3) infection, epithelial cell response/turnover associates with preserved growth and barrier function, but during undernutrition, the typical epithelial cell responses are diminished and mice demonstrate impaired growth and increased intestinal permeability [7,22]. However, the mechanism of intestinal epithelial barrier repair after *Giardia* infection remains poorly defined. Additionally, multiple pathways may determine whether barrier repair is complete or delayed after *Giardia* infection. For example, *Giardia* infection is associated with alterations in the intestinal microbiome. Animal studies show infection-associated shifts in microbial diversity and dominant bacterial taxa, including changes in Firmicutes, Bacteroidetes, and Proteobacteria [23–26]. These changes are relevant to epithelial recovery because commensal bacteria help regulate tight junction integrity, epithelial turnover, mucus production, and immune homeostasis [27–29]. During undernutrition, *Giardia* may antagonize these beneficial functions of resident microbiota [22]. Microbial metabolism also generates bioactive metabolites that influence host repair pathways, raising the possibility that *Giardia*-induced dysbiosis alters the metabolic signals needed for barrier restoration [30].

A nutrient-sensitive pathway that may connect diet, microbiota, and epithelial repair is aryl hydrocarbon receptor (AHR) signaling. Although diet can influence host resilience in simple experimental models, the intestinal microbiota may shape the epithelial responses during infection. AHR is a ligand-activated transcription factor expressed in intestinal epithelial and immune cells, where it contributes to barrier homeostasis and mucosal immunity [31–33]. Many endogenous intestinal AHR ligands are either generated through host or microbial metabolism of dietary tryptophan. They include indole-derived compounds such as kynurenine, indole-3-aldehyde, indole-3-ethanol, and indole-3-pyruvate [34–36]. These metabolites can promote epithelial repair, reduce intestinal permeability, and support mucosal healing in experimental models, with protective effects diminished when AHR signaling is disrupted [34,36–39]. Overall, these findings highlight the AHR signaling pathway as sensitive to nutrients and the microbiota, which may govern epithelial recovery following enteric infection.

Clinical evidence further supports a link between giardiasis, amino acid metabolism, and growth outcomes. In a longitudinal pediatric cohort study, *Giardia* infection was associated with reduced circulating amino acids in the plasma, including tryptophan, independent of inflammatory enteropathy, suggesting altered nutrient absorption or host-microbial metabolism rather than inflammation alone [7]. Undernourished mice similarly developed decreases in free amino acids [22]. Since tryptophan availability can modulate microbial indole production and AHR activation, reduced availability of tryptophan metabolites during giardiasis could impair epithelial repair pathways and contribute to persistent barrier dysfunction and growth impairment. We hypothesized that *Giardia*-associated disruption of tryptophan-dependent AHR signaling impairs epithelial barrier repair and contributes to associated growth failure. Using a murine model of *Giardia* infection, we tested whether dietary tryptophan availability regulates AHR-associated transcriptional responses, parasite burden, and host growth outcomes. We further asked whether tryptophan supplementation restores AHR signaling and rescues the weight defect during giardiasis independently of effects on parasite colonization.

## METHODS

### Parasites and infection

*Giardia duodenalis* clone H7 (a prototypical member of assemblage B) (ATCC# 50581) and GS-Luc-C trophozoites Kosile et al., (2026). [40] were cultured in TYI-S-33 medium at 37 °C and 5% CO2[41] until confluent before being detached on ice and washed with cold PBS prior to enumeration using a hemocytometer. Each mouse was infected by oral gavage with 1 × 10⁶ trophozoites suspended in 100 μL sterile PBS. All animal procedures were conducted in accordance with IACUC-approved protocol 2016-1128.

### Diets

The 20% protein diet (TD.91352) (RPD), isocaloric 2% protein diet (TD.92203) (LPD), 2% protein + tryptophan diet (TD.240799) (TRP), or a 2% protein + glycine diet (TD.250545) (GLY were obtained from Inotiv (Teklad™ Custom Diets, Madison, WI). The TRP (0.27% added tryptophan) and GLY (0.33% added glycine) diets were formulated by adding a comparable amount of each amino acid as is present in the 20% protein diet.

### Animals

Bartelt et al., 2013 [42] established a murine model of *Giardia* infection under conditions of dietary protein deprivation. Briefly, 3–4-week-old neonatal male C57BL/6J and BALB/c mice were purchased from The Jackson Laboratory (Bar Harbor, ME). BALB/c mice were used in these experiments because estimating parasite burden by imaging was more consistent in mice with white fur color. Mice were housed (5 mice/cage) under specific pathogen-free (SPF) conditions in ventilated cages in a room with controlled temperature and humidity and maintained on a 12-hour light/dark cycle. Animals had *ad libitum* access to food and water. Seven days before infection, mice were placed on either a standard isocaloric 20% protein diet, a 2% protein diet (low-protein diet), a 2% protein + Tryptophan diet, or a 2% protein + Glycine diet. After 7 days of dietary acclimation, mice were infected by oral gavage as described above. Uninfected control animals did not receive gavage treatment.

### Euthanasia and blood collection

At 21 days of infection, mice were anesthetized according to IACUC guidelines. Briefly, prior to retroorbital bleeding, each mouse received a Ketamine/Xylazine cocktail (100 mg/ml) administered intraperitoneally at 250 mg/kg body weight. Blood was collected using a heparinized capillary tube and was centrifuged at 5000 g at 4 °C for 10 minutes. The supernatant (plasma) was transferred to a separate tube and frozen at -80 °C prior to mass spectrometric analysis.

### Weights

Animals were weighed twice a week throughout the infection. Body weights were normalized to day 7, which served as the baseline, to account for the initial adaptation period to the experimental diets. Weight data were compiled from two or more independent infection experiments. Sample sizes for each experimental group are indicated in the corresponding figures.

### Parasite counts

Five-cm segments of the small-intestinal duodenum were excised and placed in cold PBS. Intestinal segments were opened longitudinally, minced, and suspended in 5 mL cold PBS. Trophozoites were quantified using a hemocytometer, yielding a detection limit of 2,000 trophozoites per cm of intestine. Parasite burdens were quantified at 7 and 21 days post-infection. Separate cohorts of mice were used for each time point, as intestinal tissues were collected at euthanasia. For graphical representation, samples with undetectable parasite levels were assigned the assay’s limit of detection, defined as 2,000 parasites per centimeter of intestine. All parasite burden data were subsequently log-transformed prior to analysis.

### Live bioluminescence imaging

To visualize *Giardia* (GS-Luc-C) colonization in the gastrointestinal tract, mice were administered IVISbrite D-Luciferin Potassium Salt (XenoLight) from Revvity at 150 mg/kg body weight via intraperitoneal injection. Animals were allowed to rest for 15 minutes to permit systemic distribution of the substrate. Mice were then anesthetized using 2–3% isoflurane and imaged using an IVIS imaging system (Caliper Life Sciences).

### Tissue RNA purification

Total RNA was isolated using the PureLink TRIzol™ Plus RNA Purification Kit (Invitrogen) following the manufacturer’s instructions. Briefly, 1 cm of tissue from the duodenum was homogenized in TRIzol™ to ensure complete cell lysis and RNase inactivation. After incubation, chloroform was added to induce phase separation, and samples were centrifuged to separate the aqueous RNA-containing phase from DNA and protein. The upper aqueous phase was carefully collected for downstream RNA purification. On-column DNase treatment was performed using PureLink DNase (Invitrogen) to eliminate genomic DNA contamination.

### cDNA synthesis

cDNA was synthesized using the iScript™ gDNA Clear cDNA Synthesis Kit (Bio-Rad) following the manufacturer’s instructions. Briefly, total RNA was first treated with DNase to eliminate residual genomic DNA, then heat inactivated. DNase-treated RNA was subsequently reverse transcribed using the iScript reverse transcription supermix, which contains a blend of oligo(dT) and random primers. Reactions were carried out in a thermal cycler, and the resulting cDNA was used for downstream qRT-PCR analyses.

### Targeted metabolomics

Targeted liquid chromatography-mass spectrometry (LC–MS) was used to measure indole-3-carboxaldehyde, tryptophol (indole-3-ethanol), and indole-3-pyruvic acid in mouse plasma, with indole-3-acetic-2,2-d2-acid (10 ng/mL) included as an internal standard. All standards were purchased from Sigma-Aldrich (St. Louis, MO). Calibration standards and quality controls were generated by serial dilution of mixed analyte stocks, spanning 0.065–100 ng/mL for indole-3-carboxaldehyde, 0.327–500 ng/mL for tryptophol, and 3.27–5000 ng/mL for indole-3-pyruvic acid. Plasma samples (10 µL) were extracted using an organic solvent containing 40% isopropanol, 25% methanol, and 35% water, containing 10mg/mL indole-3-acetic acid-d2 as the internal standard, followed by protein precipitation with acetonitrile and centrifugation. Chromatographic separation was performed on a Kinetex F5 column (150 × 2.1 mm) using water and acetonitrile supplemented with 0.1% formic acid. Analytes were detected by multiple reaction monitoring (MRM) with optimized transitions (Table 1). Sample order was randomized, and pooled quality controls (10 samples), standard controls, and blanks were interspersed throughout the run to monitor performance and minimize carryover. Calibration curves were linear across the tested ranges (r² > 0.99), with lower limits of quantification defined by the lowest calibration standard. Data were processed in Sciex OS software, normalized to the internal standard, and quantified by interpolation from calibration curves, with coefficients of variation ∼3-15% across analytes.

**Table 4.1.** LC–MS MRM parameters for indole metabolite analysis.

| Metabolite | Q1 (m/z) | Q3 (m/z) | Dwell Time (ms) | Role |
| --- | --- | --- | --- | --- |
| Indole-3-carboxaldehyde | 146 | 91 | 75.714 | Target Analyte |
| Tryptophol | 160 | 116 | 75.714 | Target Analyte |
| Indole-3-pyruvic acid | 204 | 130 | 75.714 | Target Analyte |
| Indole-3-acetic acid-d2 | 178 | 132 | 75.714 | Internal Standard |

### Data analysis

Sample sizes for each experimental group are reported in the corresponding figure legends, and weight data were compiled from independent infection experiments. As noted above, body weights were normalized to day 7 (diet acclimation baseline) prior to statistical testing. Longitudinal weight trajectories were analyzed in GraphPad Prism using a repeated-measures two-way ANOVA (time × group), with mixed-effects modeling (REML) applied where applicable. For parasite burden quantified by bioluminescent imaging, radiance values were analyzed using two-way ANOVA. For endpoint comparisons like mRNA levels and metabolite abundances, statistical significance was assessed using two-way ANOVA (diet × infection) with a Tukey’s multiple-comparisons test for post-hoc analysis, as indicated.

## RESULTS

### Protein malnutrition and *Giardia* infection impair weight gain and indole metabolites quantity

To investigate how giardiasis affects tryptophan metabolites and growth, we used an established murine model of giardiasis that exhibits infection- and diet-dependent growth failure [42]. Three- to four-week-old male C57BL/6J mice were fed either an RPD or an LPD for a week prior to infection with *Giardia*. Consistent with previous reports, *Giardia* infection did not impact weight gain in mice fed RPD. In contrast, infected mice fed LPD showed significant and sustained impairment in weight gain compared to uninfected controls [7,22,30,42] (Figure 1A-B).

**Figure 1:**
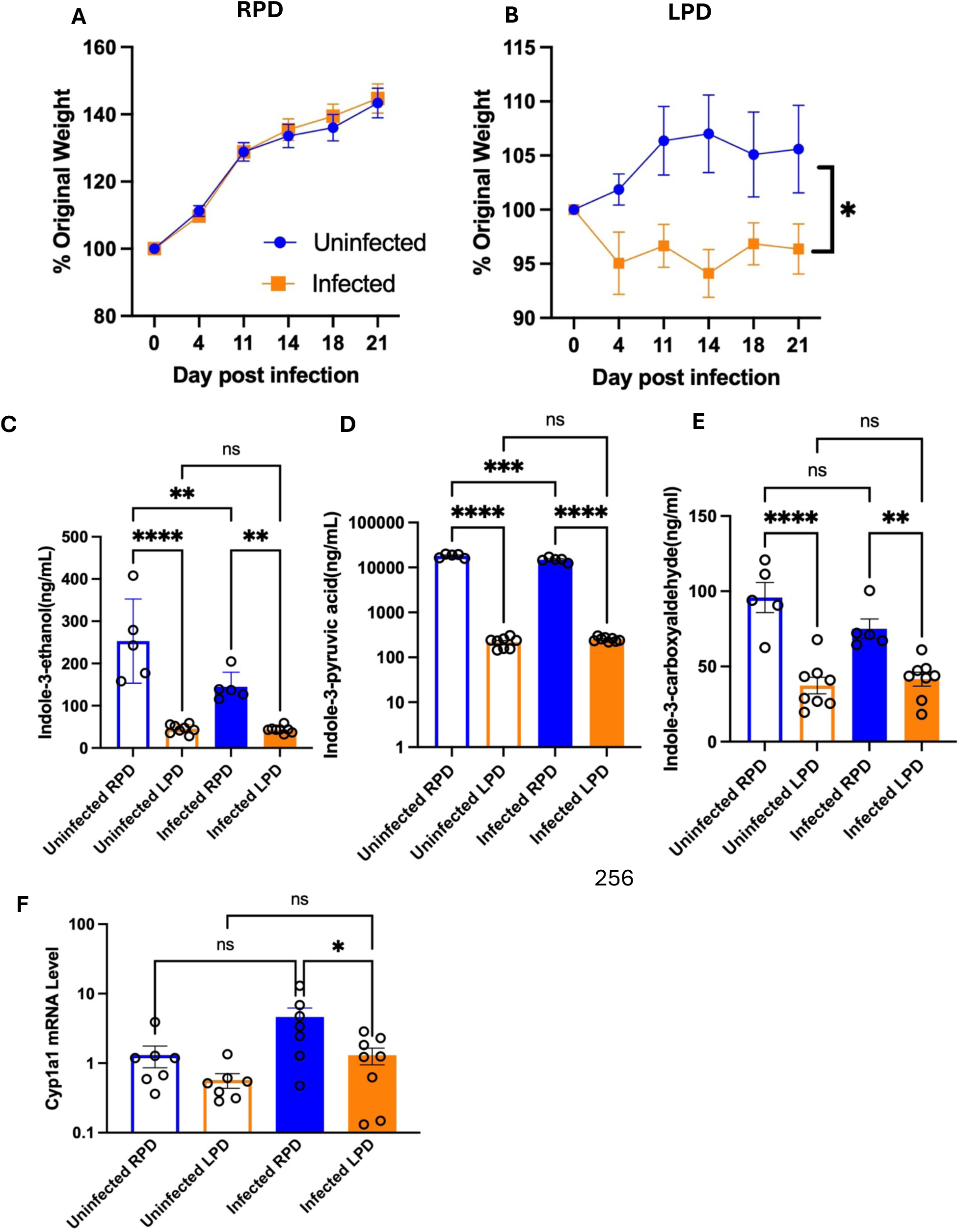
*Giardia* infection and plasma tryptophan metabolite levels in mice fed a low-protein diet (LPD) correlate to reduced weight gain. Three to four-week-old mice were maintained on RPD or LPD for 7 days and infected with *Giardia* strain GS. Mice were weighed over time and normalized to day 7 to account for dietary acclimation. (A) Weight change over time in RPD-fed mice, infected versus uninfected controls. (B) Weight change over time in LPD-fed mice, infected versus uninfected controls. (C-E) Targeted LC–MS analysis of circulating tryptophan metabolites in plasma from infected and control mice fed RPD or LPD. (F) Small intestinal Cyp1a1 mRNA levels at day 21 post-infection, measured by RT-qPCR and normalized to GAPDH. Data are presented as mean ± SEM from ≥2 independent experiments (n = 7/group). We analyzed longitudinal weight data using a two-way ANOVA with a mixed-effects model (REML). Other datasets were analyzed by Student’s *t*-test or two-way ANOVA with Tukey’s multiple-comparisons test. Statistical significance is indicated as: *P <* 0.05*, P <* 0.01*, P <* 0.001*, P <* 0.0001; ns, not significant.

Given the established role of dietary protein in shaping tryptophan availability, we next examined whether differences in protein intake influenced tryptophan metabolism and downstream aryl hydrocarbon receptor (AHR) signaling [31,35]. Targeted LC-MS analysis of plasma metabolites was performed in mice fed the LPD or RPD, with or without infection. RPD-fed mice had substantially higher plasma levels of select tryptophan-derived metabolites than LPD-fed mice, whereas infection produced only some impact relative to diet within the RPD group (Figure 1C–E). This finding suggests that protein malnutrition restricts systemic tryptophan metabolic outputs regardless of infection. Consistent with this observation, mRNA levels of Cyp1a1, a canonical transcriptional target of AHR signaling, were significantly reduced in LPD-fed mice compared with RPD-fed controls (Figure 1F). These results suggest that low dietary protein availability may limit AHR signaling pathways during *Giardia* infection.

### Tryptophan supplementation restores growth and AHR signaling independently of plasma indole abundance

To directly test whether restoring dietary tryptophan could rescue infection-related growth failure in young C57BL/6J mice, the LPD diet was modified by adding tryptophan (TRP) at a level equivalent to that in the RPD. As previously described, mice started on each diet one week before infection. As shown earlier, infected mice fed the LPD exhibited a significant growth deficit. In contrast, tryptophan supplementation prevented this infection-associated growth impairment, resulting in weight trajectories in infected mice that were comparable to those of uninfected controls. (Figure 2A–C).

**Figure 2:**
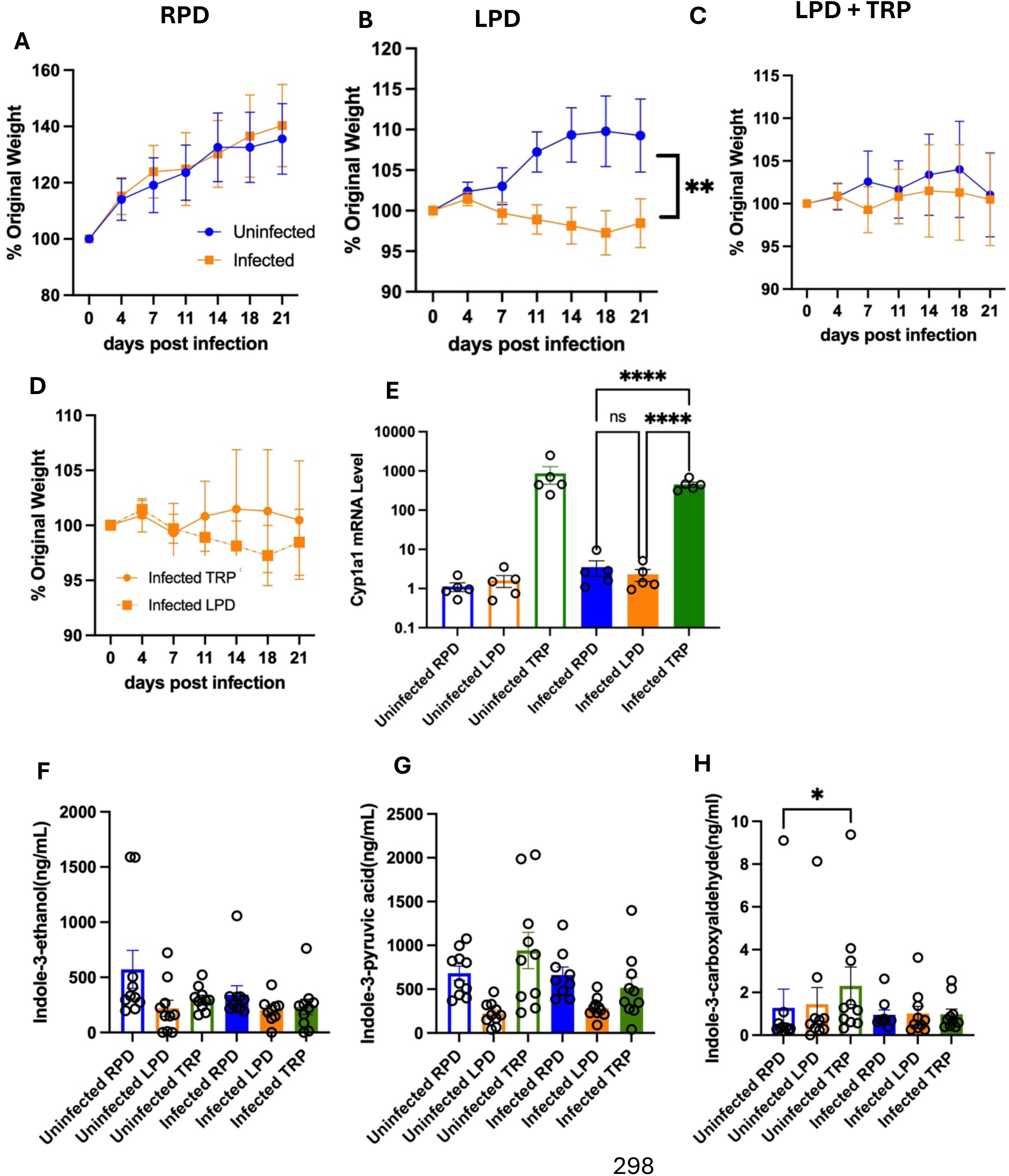
Tryptophan supplementation restores growth and enhances AHR-associated transcriptional responses without altering plasma indole levels. Three to four-week-old C57BL/6J mice were maintained on either an RPD, LPD, or TRP and either infected with *Giardia* or left uninfected as controls. Mice were weighed over time and normalized to day 7 of the diet to account for dietary acclimation. (A-C) Weight trajectories of mice fed RPD, LPD, or TRP during *Giardia* infection. (D) Merged weight change in infected mice fed TRP (C) and LPD (B). (E) Small intestinal Cyp1a1 mRNA levels in infected and uninfected mice across dietary groups, measured by RT-qPCR and normalized to GAPDH. (F - H) Targeted LC-MS quantification of plasma indole-3-carboxaldehyde, tryptophol, and indole-3-pyruvic acid in infected and uninfected mice across dietary groups. Data are presented as mean ± SEM (n = 10/group). Longitudinal weight data were analyzed using a two-way ANOVA with a mixed-effects model (REML). Other datasets were analyzed by Student’s *t*-test or two-way ANOVA with Tukey’s multiple-comparisons test. Each data point represents an individual mouse. Statistical significance is indicated as: *P <* 0.05*, P <* 0.01*, P <* 0.001*, P <* 0.0001; ns, not significant.

In parallel, tryptophan supplementation significantly increased Cyp1a1 mRNA expression in both infected and uninfected mice (Figure 2E), indicating enhanced AHR activation. Comparing infected LPD and infected TRP groups, we observed greater weight gain in TRP-fed mice than in LPD-fed mice (Figure 2D). Targeted LC-MS analysis revealed no significant differences in circulating levels of indole-3-carboxaldehyde, tryptophol, or indole-3-pyruvic acid between dietary groups after infection (Figure F-H), indicating that improved weight gain and AHR signaling occur without detectable changes in steady-state plasma indole concentrations.

### Growth preservation occurs independently of parasite burden and is reproducible in another mouse strain

To determine whether improved growth is reproducible in another strain of wild-type mice and whether parasite burdens are affected by diets using bioluminescent parasites, we fed BALB/c mice the same diets and infected them as described previously. In BALB/c mice, tryptophan supplementation again significantly improved diet-matched weight gain during infection (Figure 3A-D), demonstrating that the growth-preserving effect is consistent across genetic backgrounds. To determine whether diet altered infection intensity, parasite burdens were measured across dietary groups using bioluminescent imaging, as described in Kosile et al. (2026)[40]. Parasite burdens at 6 and 17 days post-infection were comparable across LPD and TRP dietary groups but were lower in the infected RPD group (Figure 3F), indicating that tryptophan supplementation does not measurably reduce *Giardia* intestinal colonization intensity in these animals. These data suggest the likelihood of a mechanism in which nutritional supplementation improves growth outcomes without diminishing parasite replication or enhancing parasite clearance.

**Figure 3:**
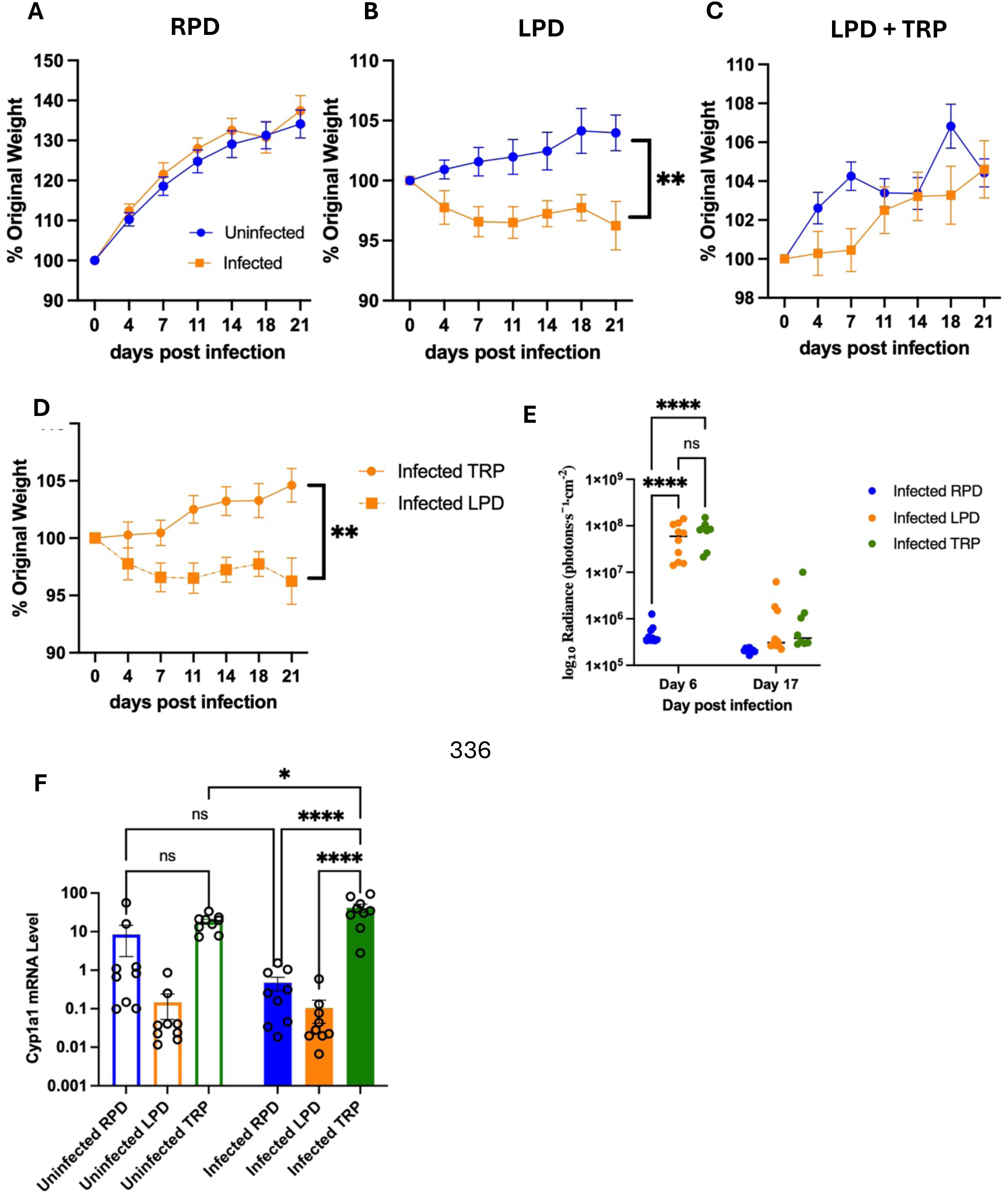
Effects of tryptophan supplementation on growth, parasite burden, and intestinal transcriptional responses in mice during BALB/c *Giardia* infection. Three-to four-week-old male BALB/c were maintained on either RPD, LPD, and TRP and either infected with *Giardia* or left uninfected as controls. Mice were weighed over time and normalized to day 7 to account for dietary acclimation and plotted as percent original weight. (A–C) Percent original weight over time for mice fed (A) RPD, (B) LPD, or (C) TRP; blue, uninfected; orange, infected. (D) Percent original weight over time for infected mice fed LPD (circle) or TRP (square). (E) Parasite burden assessed by bioluminescent imaging in infected mice fed RPD, LPD, or TRP at 6 and 17 days post-infection. (F) Small intestinal Cyp1a1 mRNA levels in uninfected and infected mice across dietary groups, measured by RT-qPCR and normalized to GAPDH. Data are presented as mean ± SEM. Longitudinal weight data were analyzed using a two-way ANOVA with a mixed-effects model (REML). Other datasets were analyzed by Student’s *t*-test or two-way ANOVA with Tukey’s multiple-comparisons test. Each data point represents an individual mouse. Statistical significance is indicated as: *P <* 0.05*, P <* 0.01*, P <* 0.001*, P <* 0.0001; ns, not significant.

Analysis of intestinal Cyp1a1 expression revealed increased expression in tryptophan-supplemented mice relative to the LPD controls (Figure 3F). This finding suggests that restoring dietary tryptophan availability is associated with increased AHR signaling activity in the intestine.

### Growth rescue is specific to tryptophan rather than general amino acid supplementation

To assess whether growth rescue reflected a general effect of amino acid supplementation, we supplemented LPD with either tryptophan (TRP) or glycine (GLY) before infection. Glycine was selected because it is not metabolized into potential Ahr ligands and is present in the RPD at levels similar to TRP. RPD-fed mice gained weight steadily regardless of infection status as previously shown (Figure 4A). Tryptophan supplementation significantly improved weight gain in infected mice, whereas glycine supplementation had minimal effect (Figure 4B–D). When we compare the infected TRP and GLY groups, we observed a significant difference in weight gain. Although the infected GLY-fed mice are slightly growth restricted compared to the TRP-fed mice, there was a significant difference between uninfected and infected GLY-fed mice (Figure 4B). Parasite burdens remained comparable across the GLY and TRP groups, while RPD-fed mice had lower early infection burdens and cleared infection faster, as shown by the radiance intensity (Figure 4E). The glycine supplementation control provides important specificity, showing that the growth improvement is not simply due to increased amino acid availability. Instead, these results indicate that growth rescue is not explained by generalized amino acid supplementation and instead points to a specific role for dietary tryptophan in shaping host physiology during infection. This specificity strengthens the rationale for focusing on tryptophan-dependent signaling and metabolism in future mechanistic studies.

**Figure 4:**
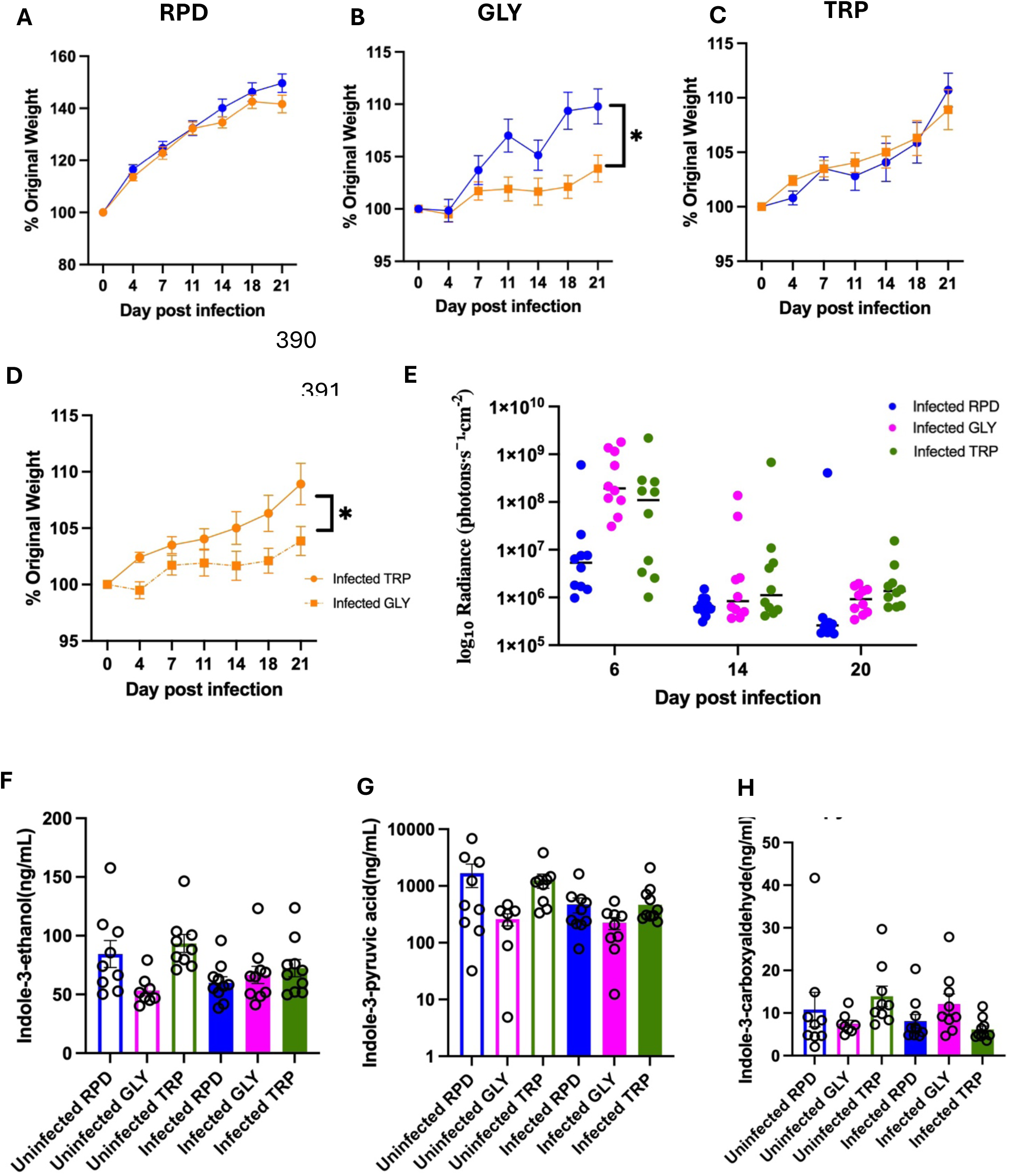
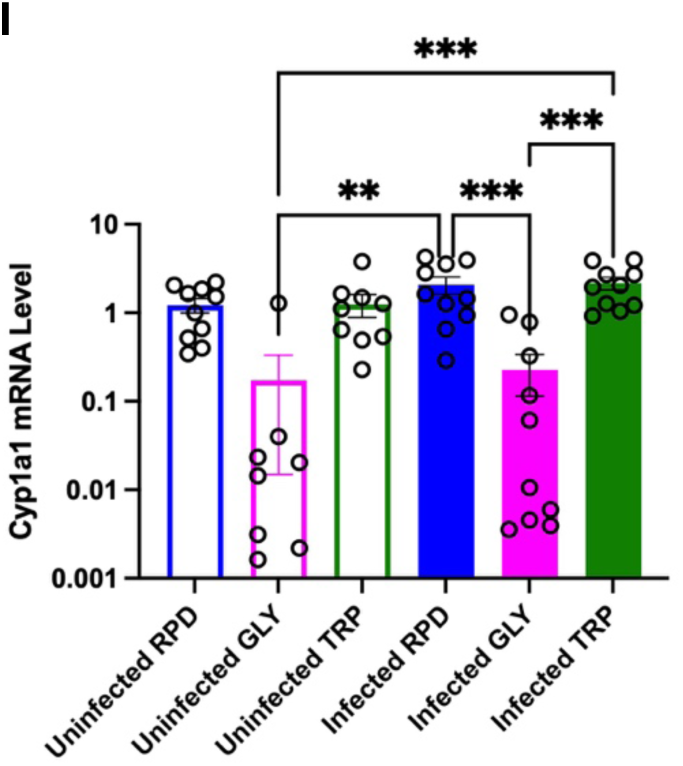
Growth rescue during giardiasis may be specific to tryptophan supplementation. Three-to-four-week-old male Balb/c mice were maintained on RPD, GLY, and TRP, or infected with *Giardia,* or left uninfected as controls, where indicated. Body weight was measured longitudinally, normalized to day 7 of the diet to account for dietary acclimation, and plotted as a percentage of the original weight. Mice were weighed over time and normalized to day 7 to account for dietary acclimation. (A) Percent original weight over time in RPD-fed mice, uninfected versus infected. (B–C) Percent original weight over time in infected mice fed (B) TRP or (C) GLY. (D) Direct comparison of percent original weight over time in infected mice fed TRP versus GLY. (E) Small intestinal parasite burden assessed at 6, 14, and 20 days post-infection across dietary groups by BLI. (F-H) Targeted LC-MS quantification of plasma indole-3-carboxaldehyde, tryptophol, and indole-3-pyruvic acid in infected and uninfected mice across dietary groups. (I) Small intestinal Cyp1a1 mRNA levels in infected and uninfected mice across dietary groups, measured by RT–qPCR and normalized to GAPDH. Data are presented as mean ± SEM (n = 10/group). Longitudinal weight data were analyzed using a two-way ANOVA with a mixed-effects model (REML). Other datasets were analyzed by Student’s *t*-test or two-way ANOVA with Tukey’s multiple-comparisons test. Each data point represents an individual mouse. Statistical significance is indicated as: *P <* 0.05*, P <* 0.01*, P <* 0.001*, P <* 0.0001; ns, not significant.

Analysis of intestinal Cyp1a1 expression revealed an increase in tryptophan-supplemented mice, whereas glycine supplementation did not significantly alter Cyp1a1 levels relative to LPD controls, as observed in previous experiments (Figure 4I). The infected RPD-fed mice exhibited similarly elevated Cyp1a1 expression. These results demonstrate that increased Cyp1a1 expression matches the growth-rescuing effect of tryptophan supplementation and further support the specificity of tryptophan-dependent AHR signaling responses in shaping host growth outcomes during giardiasis. Additional targeted LC-MS to quantify levels of IET, IPYA, and I3A showed that across groups, circulating indole metabolite levels were variable and most pairwise comparisons were not statistically significant (Figure 4F-H). The observed pattern of plasma indole levels did not consistently match the growth-rescue phenotype. This suggests that circulating indole levels may not fully reflect the local intestinal AHR ligand availability that regulates tryptophan-dependent AHR signals during *Giardia* infection and may not reflect.

## DISCUSSION

This study extends findings that dietary protein plays a crucial role in determining host growth outcomes during *Giardia duodenalis* infection to additional *Giardia* strains, hosts, and laboratory environments, through host-driven mechanisms that are independent of the parasite burden. We tested GS/GS-Luc parasites in C57BL/6J and BALB/c mouse models and found that, consistent with previous H3 cyst infection studies, mice maintained on LPD exhibit a weight defect, whereas those on RPD maintain growth despite infection[42]. We identified tryptophan-dependent host signaling as a specific nutrient affecting infection-related growth failure. *Giardia* infection on LPD reduced weight gain and reduced expression of the aryl hydrocarbon receptor (AHR)– target gene Cyp1a1. Conversely, mice on a nutritionally complete (RPD) diet maintained consistent growth despite infection. Importantly, supplementing LPD with dietary tryptophan rescued growth failure to diet-matched controls across different mouse strains and enhanced AHR signaling without decreasing intestinal parasite numbers, emphasizing host nutritional status as a key factor in disease outcomes. Our findings support a model in which nutrient availability affects epithelial repair and growth during giardiasis through host-driven metabolic and signaling pathways, rather than by influencing parasite burden[7].

The selective emergence of growth failure in mice fed LPD is consistent with epidemiological observations linking giardiasis to growth stunting in malnourished children in low-and middle-income countries, where infection is often chronic and subclinical rather than acutely diarrheal [3,43,44]. Experimental studies similarly demonstrate that feeding mice a low-protein diet (LPD) exacerbates growth impairment and intestinal dysfunction during enteric infections [42]. Consistent with prior evidence, we show that *Giardia* infection impaired growth in mice maintained on LPD, whereas infected mice fed a regular-protein diet (RPD) maintained normal growth patterns[7,30]. These findings indicate that parasite exposure alone is insufficient to drive growth failure and instead highlight host nutritional status as a key determinant of disease severity.

Dietary protein intake is a major determinant of systemic and intestinal tryptophan availability, with protein restriction reducing circulating and tissue tryptophan pools [45–47]. Tryptophan metabolites are well-established regulators of Aryl hydrocarbon receptor (AHR), linking diet and microbial metabolism to host signaling pathways [31,35]. In barrier tissues such as the intestine, skin, and lungs, AHR signaling plays a crucial role in maintaining epithelial homeostasis, supporting tissue adaptation to changing conditions, and supporting tissues against environmental stressors (e.g., dietary compounds, microbial metabolites, pollutants, or pathogens) [48,49]. Conversely, microbiota can also convert tryptophan into barrier-toxic indoles, like indoxyl sulfate, that accumulate in urinary profiles of undernourished mice with *Giardia* infection[30]. In this study, reduced Cyp1a1 expression in LPD–fed mice and its restoration following tryptophan supplementation suggest that dietary tryptophan availability beneficially modulates AHR signaling in the intestine during giardiasis. Notably, increased Cyp1a1 expression consistently tracked with improved growth outcomes across mouse strains and dietary interventions. These findings align with prior reports showing that AHR signaling can support epithelial integrity and tissue adaptation independently of classical inflammatory cytokine pathways [49].

A key finding of this study is the temporal dissociation between parasite burden and weight around day 14 to 20 post-infection. Although parasite loads were higher in LPD-fed mice during the early stages of infection, intestinal parasite burdens were comparable between dietary groups by the end of the infection period, and RPD-fed mice cleared parasites more quickly[42]. Despite similar parasite loads during this period, significant differences in weight gain and Cyp1a1 responses remained between groups. This suggests that variations in growth outcomes cannot be fully attributed to differences in parasite burden during the later stage of infection. Our findings are consistent with the broader concept that *Giardia* pathogenicity is not solely determined by colonization but by host-specific factors that influence tissue repair, tolerance, and physiological adaptation. This also mirrors clinical observations in which some individuals may experience prolonged diarrhea, while others may harbor *Giardia* asymptomatically, and regardless of symptoms some may experience severe growth impairment and intestinal dysfunction [50,51]. Our data, therefore, suggest that host nutritional status and epithelial signaling capacity contribute to growth impairment during giardiasis, particularly as infection resolves.

Although dietary tryptophan supplementation rescued the growth defect and increased Cyp1a1 expression, these effects were not accompanied by consistent changes in the circulating indole metabolites measured in the plasma. The plasma levels of the indoles measured above (figures 1c-e) varied across groups, but the overall pattern did not match the growth-preservation phenotype (figure 4). However, we see a pattern of reduced circulating IPYA in plasma of infected animals on RPD and TRP, which may indicate increased turnover or utilization of these metabolites in the gut during infection. This may also suggest that plasma indole levels do not fully capture the biologically relevant tryptophan-dependent signals during giardiasis. AHR ligands may act locally within the intestinal lumen, undergo turnover rapidly, or differ between the lumen, tissue, and plasma. Similar discrepancies between systemic metabolite abundance and tissue-specific signaling have been reported previously, particularly for AHR ligands that act locally at barrier sites and turn over rapidly [32,35,36]. Previous studies also indicated that tryptophan metabolite levels may vary across fecal, intestinal, and circulating compartments such as plasma. These studies also showed that fecal as well as circulating tryptophan metabolite measurements closely reflect intestinal AHR activity [52,53].

These findings suggest that local intestinal tryptophan metabolism or epithelial sensitivity to available ligands, rather than steady-state plasma concentrations, may determine AHR signaling during giardiasis. Intestinal Cyp1a1 expression may provide a more useful readout for AHR-associated signaling than the level of circulating indoles in the plasma. Future studies would consider measuring tryptophan metabolites in intestinal tissue, luminal content, or fecal samples to accurately determine whether local metabolite availability correlate with AHR activation and growth-rescue.

We used glycine supplementation as a control to test whether the growth rescue observed with tryptophan reflects a general amino acid effect. Although the glycine cohort did not include matched LPD controls (limiting cross-cohort comparisons), glycine did not recapitulate the TRP-associated phenotype within the same experiment: TRP outperformed GLY for weight gain during infection and was associated with increased intestinal Cyp1a1 expression. This shows that the observed phenotype is not due to a random amino acid in the LPD. This specificity aligns with previous research indicating that tryptophan selectively engages pathways involved in immune modulation and epithelial function, including AHR signaling, whereas glycine supplementation did not recapitulate these effects [46,54]. Because many biologically active tryptophan metabolites are generated through microbial metabolism, these findings also support a model in which dietary tryptophan availability influences host responses indirectly through microbiome-dependent AHR ligand production. These data support the conclusion that dietary tryptophan plays a distinct role in shaping host responses during *Giardia* infection. Notably, tryptophan supplementation did not increase weight gain in uninfected mice, particularly in the C57BL/6J mice suggesting that its effects are context-dependent and primarily act to restore growth under conditions of nutritional or infection-associated stress rather than promoting baseline growth. Together, these data support the conclusion that dietary tryptophan plays a distinct role in shaping host responses during *Giardia* infection.

This work identifies dietary tryptophan availability as a key host factor that determines growth outcomes during giardiasis independently of measurable reductions in parasite burden. By linking nutritional status to epithelial-intrinsic AHR signaling, these findings provide a mechanistic framework for understanding why *Giardia* infection disproportionately affects growth in malnourished populations. Targeting nutrient-sensitive host pathways may therefore represent a complementary strategy for mitigating the long-term consequences of giardiasis in settings where malnutrition is prevalent.

## Acknowledgments

This work was supported by the Georgetown University Division of Comparative Medicine and the Small Animal Imaging Shared Resource and the Metabolomics Core Facility of the Lombardi Comprehensive Cancer Center.

